# No observed effect concentration (NOEC) and minimal effective dose (MED): estimation of non-experimental doses

**DOI:** 10.1101/2023.08.23.554562

**Authors:** Ludwig A. Hothorn

## Abstract

In in-vitro or in-vivo bioassays, the no observed effect concentration (NOEC) is often determined. This simple procedure has several disadvantages, including the limitation of being able to estimate only experimental doses. Linear interpolation between adjacent doses overcomes this drawback while maintaining the level of a familywise error rate (FWER) using multiple contrast tests.

## 1 Introduction

Both minimum effective dose and no observed effect concentration are related approaches, whereas NOEC=MED-1 within a proof of hazard approach. Recently, the concept of MED was used by estimating a MED of 50% in a pain reduction study [7], see Figure 1.

**Figure 1:**
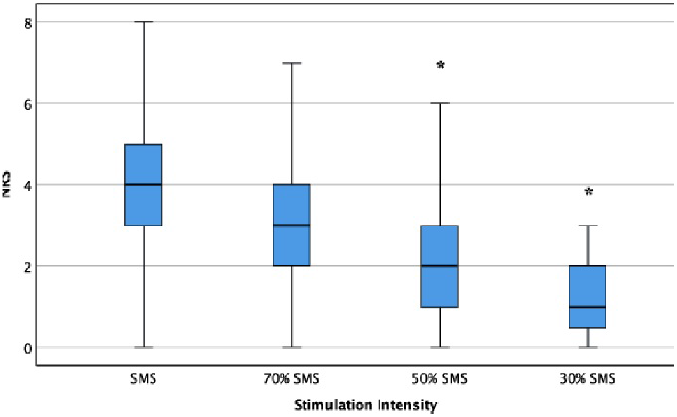
Estimated *MED* = 50% for pain reduction by stimulation.

**Figure 2:**
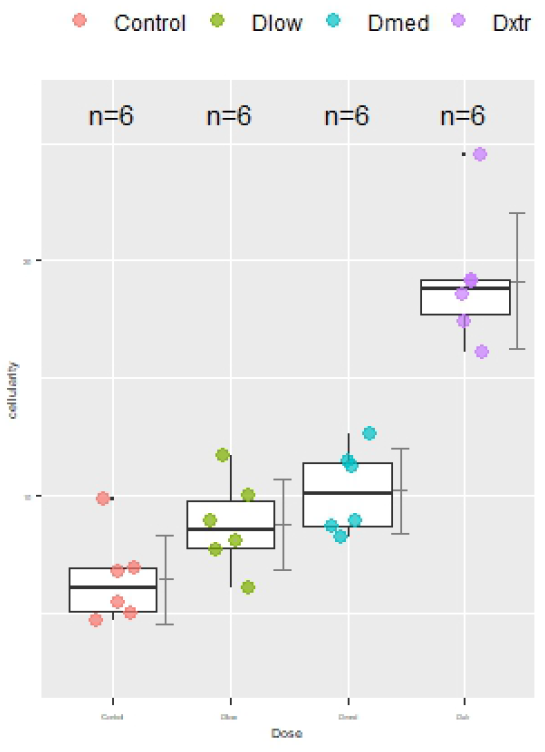
BALB cellularity data

However, this concept has several and serious drawbacks. E.g., the MED depends strongly on the design, particularly on sample sizes *n*_*i*_, on variance (and possible variance heterogeneity), and on the number of doses (together with a negative control in the one-way layout). Furthermore, only experimental doses can be estimated as MED, so in the above example the MED can only take the values of 70, 50, 30% . Especially this is a substantial disadvantage compared to alternative concepts like the benchmark dose in risk assessment [8]. The disadvantage of the first problem cluster can be partially limited by following the recommended design in related guidelines. The OECD draft guidance for field trials with agrochemicals [1] propose a design with 4 concentrations and samples sizes of *n*_*i*_ = 6. The second disadvantage can be limited by linear interpolation between the doses *D*_*i*_ between MED and MED-1 the focus of this paper.

A simple linear interpolation between adjacent doses can be used: *D*_*interpolated*_ = (1*−η*)*D*_*MED−*1_+ *ηD*_*MED*_ proposed by [3]. However, since the value of *η* is unknown a-priori, and indeed data-dependent, one can span a grid of e.g. 9 *η* values (*>* 0; *<* 1). This grid can be models as multiple contrasts, in addition to the contrast of the Dunnett procedure [4]-commonly used for MED estimation. This greatly increased multiplicity. But its inherent conservatism is minimized by the high correlation between the (*k* + 9) contrasts.

## 2 An interpolated MED approach

The MED can be simply defined as *MED* = argmin_*i*:1,…,*k*_ : *μ*_*i*_ *> μ*_0_ + Δ [9], [2]) with a threshold of Δ = 0. An unbiased estimation can be achieved by the one-sided Dunnett procedure for the assumption of *N* (*μ*_*i*_, *sigma*^2^) or with modified variances estimator or adjusted df’s for the more realistic assumption of *N* 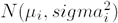 [6]. The Dunnett procedure can be formulated as maximum contrast test *t*_*MCT*_ = *max*(*t*_1_, …, *t*_Ξ_) based on the *ξ* marginal contrasts 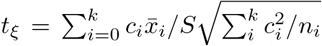 where the contrast coefficient determines the specific kind of multiple contrast test. For a simple balanced design [*C, D*_1_, *D*_2_, *D*_3_] the contrast matrix for one-sided comparisons versus a control are:

Assuming *D*_2_ was estimated as an experimental MED, the following 9 marginal contrasts are added to the matrix to estimate the interpolated MED:

The interpolated MED is then 0.7*D*_1_ + 0.3*D*_2_ if the smallest adjusted p-value occurs at contrast *c*_*f*_.

## 3 Example and R-code

As example the endpoint cellularity in a lymph node assay is considered in a balanced one-way layout with a control and three doses (assumed as 0, 10, 50, 200 mg) [5] (see the raw data in its table 1):

**Table 1:** Contrast matrix Dunnett test k=3+1.

| $c_i$ | C | $D_1$ | $D_2$ | $D_3$ |
| --- | --- | --- | --- | --- |
| $c_a$ | -1 | 0 | 0 | 1 |
| $c_b$ | -1 | 0 | 1 | 0 |
| $c_c$ | -1 | 1 | 0 | 0 |

**Table 2:** Contrast matrix to estimate interpolated MED for a k=3+1 design.

| $c_i$ | C | $D_1$ | $D_2$ | $D_3$ |
| --- | --- | --- | --- | --- |
| $c_a$ | -1 | 0 | 0 | 1 |
| $c_b$ | -1 | 0 | 1 | 0 |
| $c_c$ | -1 | 1 | 0 | 0 |
| $c_d$ | -1 | 0.9 | 0.1 | 0 |
| $c_e$ | -1 | 0.8 | 0.2 | 0 |
| $c_f$ | -1 | 0.7 | 0.3 | 0 |
| $c_g$ | -1 | 0.6 | 0.4 | 0 |
| $c_h$ | -1 | 0.5 | 0.5 | 0 |
| $c_i$ | -1 | 0.4 | 0.6 | 0 |
| $c_j$ | -1 | 0.3 | 0.7 | 0 |
| $c_k$ | -1 | 0.2 | 0.8 | 0 |
| $c_l$ | -1 | 0.1 | 0.9 | 0 |

**Table 3:** Dunnett test of cellularity data example.

| Contrast | estimate | se | test stat | p-value |
| --- | --- | --- | --- | --- |
| 10 - Control | 2.32 | 1.22 | 1.90 | 0.0884 |
| 50 - Control | 3.76 | 1.17 | 3.20 | **0.0061** |
| 200 - Control | 12.68 | 1.54 | 8.22 | 0.0000 |

**Table 4:** Contrast matrix for interpolated MED test for cellularity data example.

| Contrast | Control | 10 mg | 50 mg | 200 mg |
| --- | --- | --- | --- | --- |
| 10 - Control | -1 | 1 | 0 | 0 |
| 50 - Control | -1 | 0 | 1 | 0 |
| 200 - Control | -1 | 0 | 0 | 1 |
| weight01 | -1 | 0.10 | 0.90 | 0 |
| weight02 | -1 | 0.20 | 0.80 | 0 |
| weight03 | -1 | 0.30 | 0.70 | 0 |
| weight04 | -1 | 0.40 | 0.60 | 0 |
| weight05 | -1 | 0.50 | 0.50 | 0 |
| weight06 | -1 | 0.60 | 0.40 | 0 |
| weight07 | -1 | 0.70 | 0.30 | 0 |
| weight08 | -1 | 0.80 | 0.20 | 0 |
| weight09 | -1 | 0.90 | 0.10 | 0 |

**Table 5:** Interpolated MED test for cellularity data example.

| Contrast | estimate | se | test stat | p-value |
| --- | --- | --- | --- | --- |
| 10- Control | 2.32 | 1.22 | 1.90 | 0.0920 |
| 50 - Control | 3.76 | 1.17 | 3.20 | 0.0067 |
| 10 - Control | 12.68 | 1.54 | 8.22 | 0.0000 |
| 0.1/0.9 | 3.61 | 1.12 | 3.22 | 0.0065 |
| 0.2/0.8 | 3.47 | 1.08 | 3.21 | 0.0067 |
| 0.3/0.7 | 3.33 | 1.05 | 3.16 | 0.0074 |
| 0.4/0.6 | 3.18 | 1.04 | 3.06 | 0.0092 |
| 0.5/0.5 | 3.04 | 1.04 | 2.93 | 0.0121 |
| 0.6/0.4 | 2.89 | 1.05 | 2.76 | 0.0176 |
| 0.7/0.3 | 2.75 | 1.07 | 2.56 | 0.0265 |
| 0.8/0.2 | 2.61 | 1.11 | 2.35 | **0.0406** |
| 0.9/0.1 | 2.46 | 1.16 | 2.12 | 0.0619 |

The experimental MED was estimated to 50 mg by means of the R-code:

`library(multcomp); library(sandwich)`

`mod1<-lm(cell_BALB∼group, data=vor)`

`eMED<-summary(glht(mod1, linfct = mcp(group = “Dunnett”)`,

`vcov=vcovHC, alternative=“greater”))`

To estimate the interpolated the following contrast matrix *xdl* is used:

The interpolated MED was estimated by means of the R-code:

`summary(glht(mod1, linfct = mcp(group =xdl),vcov=vcovHC,alternative=“greater”))`

The interpolated MED is then 0.8 *\* D*_1_ = 10 + 0.2*\* D*_2_ = 50 = 18*mg* instead of 50 mg in the common used approach. The multiplicity price seems to be acceptable: the adjusted p-value decreases for contrast 0.8*/*0.2 from 0.041 to 0.039 if we would a-priori know the true interpolated MED

## 4 Summary

Using a grid of multiple contrasts, linear interpolation between adjacent doses can be performed to estimate an interpolated NOEC, respective MED, with a modified Dunnett test. The related R code is straightforward. For risk assessment the no observed effect concentration (NOEC) can be estimated accordingly within the framework of proof of hazard. Comparison to the benchmark dose approach is of interest for future research. The approach presented here needs very few assumptions: i) a simple MED definition, ii) point-zero hypothesis tests, iii) control of the FWER, and iv)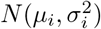, knowing that the Dunnett test is relatively robust to related data-specific violations.

